# Mena and VASP are required for vascular smooth muscle relaxation

**DOI:** 10.1101/2023.04.21.537622

**Authors:** Lea Ulrich, Carla Gliem, Dieter Groneberg, Timo Frömel, Kai Schuh, Ingrid Fleming, Peter M. Benz

## Abstract

Enabled/vasodilator stimulated phosphoproteins (Ena/VASP) proteins are important regulators of the cytoskeleton, linking kinase signaling pathways to actin assembly. In mammals, the Ena/VASP family of proteins consists of mammalian enabled (Mena), VASP, and Ena-VASP-like protein (EVL). The proteins are well known targets of cAMP- and cGMP-dependent protein kinases, PKA and PKG, respectively. Given the importance of cyclic nucleotide signaling in mediating vasodilation, we investigated the role of Ena/VASP protein in vascular smooth muscle relaxation. Whereas VASP and Mena were strongly expressed in vascular smooth muscle cells, EVL was undetectable in the arterial wall and EVL-deficiency had no impact on agonist-induced smooth muscle relaxation. VASP deletion impaired the acetylcholine (ACh)- and nitric oxide (NO)-induced relaxation murine mesenteric arteries *ex vivo*. Similarly, the ACh-induced and NO-dependent relaxation of aorta from 7-month-old but not 3- month-old VASP^-/-^ mice was also reduced. Aortas from animals lacking VASP and expressing only minimal amounts of Mena displayed significantly impaired relaxations in response to NO, cAMP and cGMP stimulation. These results suggest that Mena and VASP play an important role in agonist induced smooth muscle relaxation and functionally compensate for each other.

## Introduction

Actin binding proteins are of crucial importance for the spatiotemporal regulation of actin dynamics, thereby mediating a tremendous range of cellular processes. The enabled/vasodilator-stimulated phosphoprotein (Ena/VASP) family is one of the most fascinating and versatile family of actin regulating proteins. The proteins are localized at sites of high actin turnover, including focal adhesions, the leading edge of lamellipodia, and the tips of filopodia, where they promote actin polymerization and cell migration (Faix and Rottner, 2022). In mammals, the Ena/VASP family of proteins consists of mammalian enabled (Mena), VASP, and Ena-VASP-like protein (EVL). The family members share a tripartite domain organization of an N-terminal Ena/VASP homology 1 (EVH1) domain, a central proline-rich region (PRR), and an EVH2 domain at the C terminus (Figure 1) and (Sechi and Wehland, 2004). The mammalian Ena/VASP proteins are well known substrates of serine/threonine kinases, and human VASP harbors the three phosphorylation sites serine 157 (S157), serine 239 (S239), and threonine 278 (T278) (Benz et al., 2009). Whereas the first two phosphorylation sites are conserved in Mena, EVL contains only the first site. In cells, S157 and S239 in VASP are phosphorylated by cAMP-dependent protein kinase (PKA) in that order of preference. Conversely, cGMP-dependent protein kinase (PKG), preferentially phosphorylates S239 before S157. More recently, the AMP-activated protein kinase (AMPK) was reported to phosphorylate T278 (Benz et al., 2009; Blume et al., 2007). Functionally, S157 phosphorylation has been shown to regulate the subcellular distribution and the SH3 domain mediated interactions of VASP. In contrast, S239 and T278 phosphorylation synergistically impaired Ena/VASP-driven actin filament assembly *in vitro* and in living cells (Benz et al., 2008; Benz et al., 2009).

**Figure 1.**
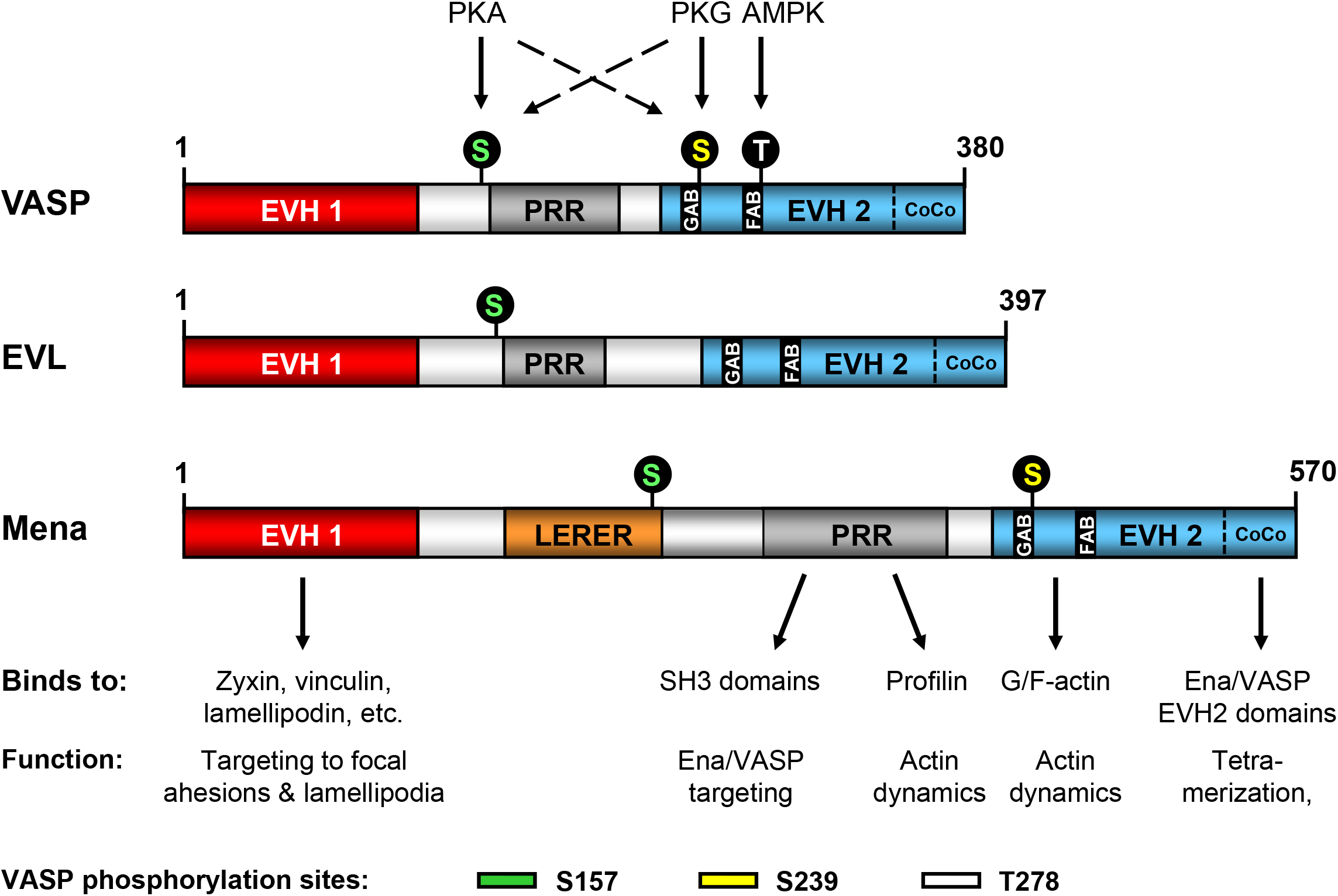
Domain organization and phosphorylation sites of Ena/VASP proteins along with their binding partners and associated functions. EVH1: Ena/VASP homology 1, PRR: proline-rich region, LERER: low complexity region harbouring LERER repeats (unique to Mena), EVH2: Ena/VASP homology 2, GAB: G-actin binding site, FAB: F-actin binding site, CoCo: coiled coil motif required for tetramerization. Numbering according to the predominant human protein isoforms. Serine and threonine phosphorylation sites and the respective kinases are also indicated. In contrast to VASP S239 (which is present in Mena but not in EVL) and T278 (which is unique to VASP), the S157 phosphorylation site is structurally/functionally conserved in all Ena/VASP proteins (color-coded circles).

Regulation of smooth muscle contractility is essential for many important biological processes including blood pressure control (Schlossmann et al., 2003). In principle, smooth muscle contraction can be induced by an increase in the cytosolic concentration of Ca^2+^ and/or the increased sensitivity of the contractile apparatus towards Ca^2+^ (Morgado et al., 2012; Schlossmann et al., 2003). Relaxation of smooth muscle is mediated by the NO/cGMP/PKG and cAMP/PKA signaling pathway, which lowers the cytosolic Ca^2+^ level and/or reduces the calcium sensitization of the contractile elements. However, phosphorylation by PKA and PKG could also modulate the cytoskeletal architecture of smooth muscle cells (Schlossmann et al., 2003; Tang, 2015; Yamin and Morgan, 2012). Given that VASP is highly expressed in smooth muscle cells and a well characterized PKA and PKG substrate, it is tempting to speculate a role for VASP in cAMP- and cGMP-mediated smooth muscle relaxation. Surprisingly, however, the cAMP- and cGMP-induced relaxation of VASP-deficient vascular smooth muscle was reported to be undistinguishable from that observed in wild-type controls. Functional redundancy of other Ena/VASP proteins was suggested to be the reason for the latter observation (Aszodi et al., 1999). In the present study, we readdressed the role of Ena/VASP proteins for smooth muscle relaxation and addressed a potential functional redundancy of the proteins.

## Results

Vascular (aortic) smooth muscle relaxation was addressed previously in tissues from 129sv inbred mice carrying a mutant VASP allele (Aszodi et al., 1999). To test a potential impact of the genetic background, we assessed the agonist-induced relaxation of vascular smooth muscle from VASP-deficient mice (Hauser et al., 1999) that had been back-crossed to C57BL/6J background for more than 12 generations (Benz et al., 2008). Thoracic aortas from 3-month-old wild-type or VASP^-/-^ mice were cut into rings and mounted in a myograph in the presence of diclofenac (10 μM). After equilibration (at least 60 minutes at 37°C), rings were precontracted with phenylephrine (1 μM). To investigate endothelial nitric oxide synthase (eNOS)-dependent smooth muscle relaxation, rings were treated with increasing concentrations of acetylcholine (ACh). eNOS-independent relaxation was tested in presence of the pharmacologic eNOS inhibitor N-nitro-L-arginine methyl ester (L-NAME, 300 μM) either using increasing concentrations of the NO-donor; dimethylamine nonoate (DEA-NONOate), or the stable cGMP analog; 8-Br-pCPT-cGMP (cGMP), to directly activate PKG. Responses to direct PKA activation were determined by monitoring responses to the stable cAMP analog; Sp-5,6-DCl-cBIMPS (cAMP). Consistent with the previous report (Aszodi et al., 1999), VASP-deficiency had no impact on the relaxation of aortic rings from 3-month-old animals (Figure 2G-J). However, when aortic rings from 7-month-old animals were studied, ACh- and NO-induced smooth muscle relaxation was significantly impaired in VASP-deficient vs. wild-type mice (Figure 2K and L). The relaxations induced by cGMP and cAMP, on the other hand, were comparable between the two strains (Figure 2 M and N). The aorta is a large conductance vessel that does not contribute to the regulation of blood pressure. Therefore, we assessed vascular reactivity in resistance arteries. Murine mesenteric arteries expressed VASP in endothelial and smooth muscle cells (Figure 3A-F). Similar to the results obtained in aortic rings from aged mice, ACh- and NO-induced smooth muscle relaxation was significantly impaired in 2^nd^ order mesenteric arteries from 4 moth old VASP-deficient animals. Again, the relaxation elicited by the cGMP analog was again unchanged compared to arteries from wild-type mice (Figure 3 G-I).

**Figure 2.**
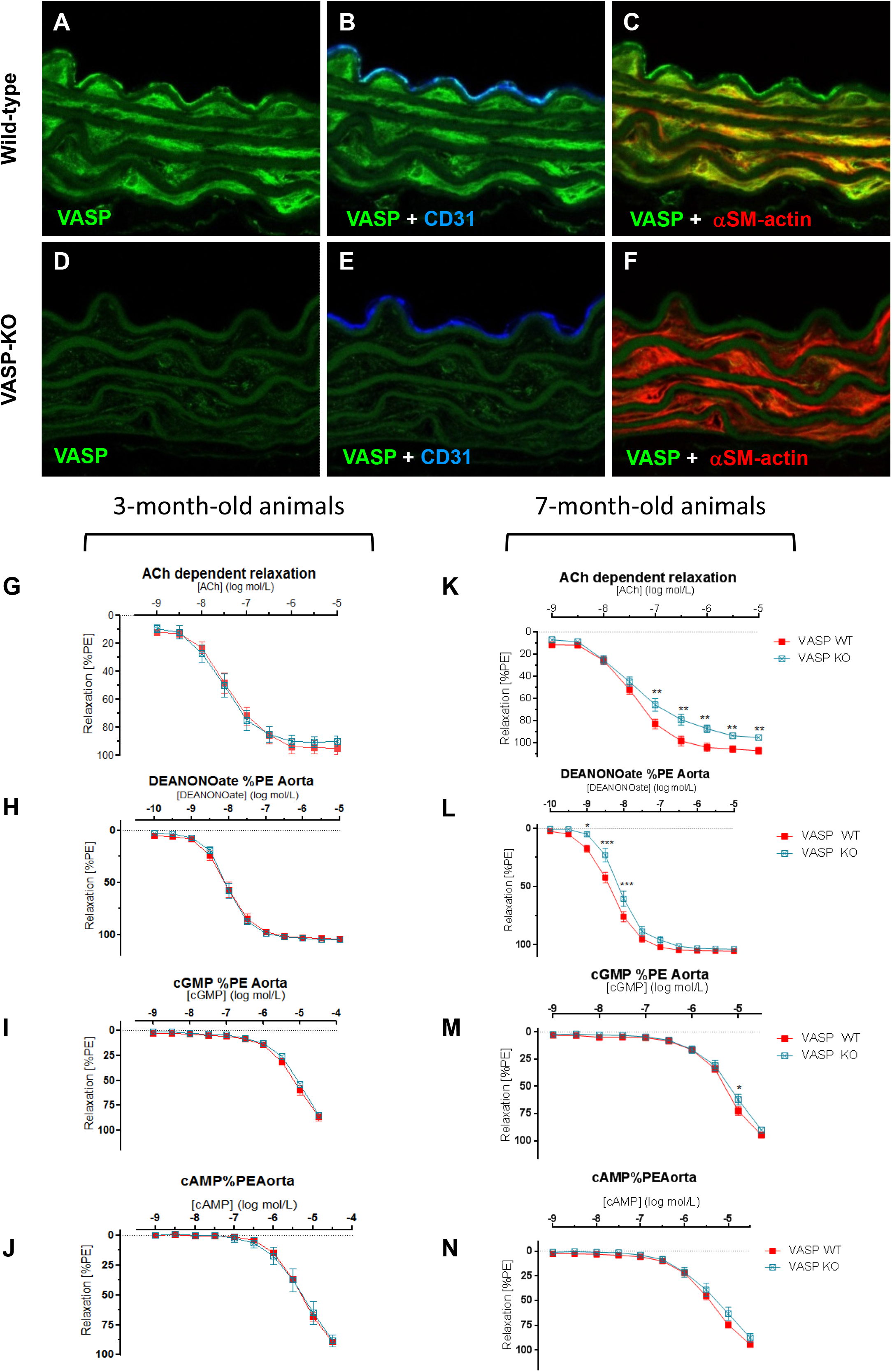
Acetylcholine- and NO-dependent relaxation of VASP-deficient aortic rings is impaired in 7-month-old animals. (**A-F**) Confocal microscopy demonstrating expression of VASP in endothelial cells and smooth muscle cells of wild-type but not VASP^-/-^ mouse aorta. Staining with CD31- and α-smooth muscle actin-specific antibodies was used to identify endothelial cells and smooth muscle cells, respectively. Representative images from 4 independent experiments are shown. (**G-N**) Myograph experiments with aortic rings from 3-month-old (G-J, N=5 animals, 4 rings per animal) and 7-month-old (K-N, N=8 animals, 4 rings per animal) VASP^-/-^ mice or wild-type controls. Aortic rings were pre-contracted with 1 μM phenylephrin (PE) and then relaxed with increasing concentrations of acetylcholine (ACh), the NO-donor DeaNONOate, or the cAMP- or cGMP-analogs Sp-5-6-DCI-BIMPS (cAMP) and 8-Br-pCPT-cGMP (cGMP), respectively. Data represent SEM ± SD, two-way ANOVA, *p < 0.05; **p < 0.01; ***p < 0.001

**Figure 3.**
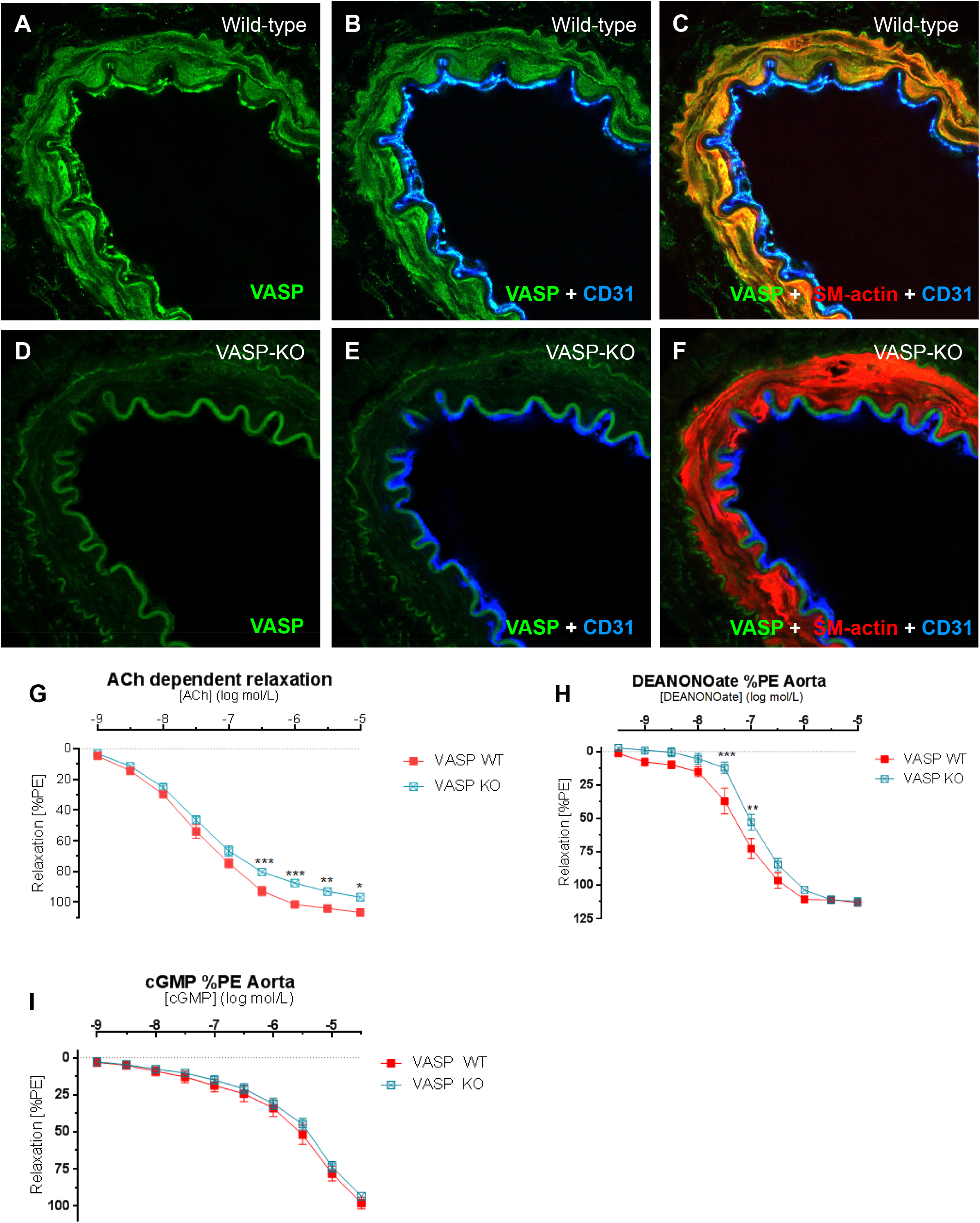
Acetylcholine- and NO-dependent relaxation of VASP-deficient mesenteric artery rings is impaired. (**A-F**) Confocal microscopy demonstrating expression of VASP in endothelial cells and smooth muscle cells of wild-type but not VASP^-/-^ mouse mesenteric artery. Staining with CD31- and α-smooth muscle actin-specific antibodies was used to identify endothelial cells and smooth muscle cells, respectively. Representative images from 4 independent experiments are shown. (**G-I**) Myograph experiments with 2^nd^ order mesenteric artery rings from 4-month-old VASP^-/-^ mice or wild-type controls (N=6 animals, 4 rings per animal). Aortic rings were pre-contracted with 1 μM phenylephrin (PE) and then relaxed with increasing concentrations of acetylcholine (ACh), the NO-donor DeaNONOate, or the cGMP-analog 8-Br-pCPT-cGMP (cGMP), respectively. Data represent SEM ± SD, two-way ANOVA, *p < 0.05; **p < 0.01; ***p < 0.001.

Next, we investigated EVL protein expression of in the wall of the mouse aorta. Although we have previously reported EVL expression in endothelial cells and blood vessels of the mouse retina (Zink et al., 2021), we could not detect the presence of EVL protein in either the smooth muscle or endothelial cells of the aorta (Figure 4A-F). Consistent with low/undetectable EVL expression in smooth muscle cells of the aorta, EVL-deficiency had no impact on the relaxation of aortic rings in response to ACh, NO, cGMP or cAMP stimulation (Figure 4 G-J).

**Figure 4.**
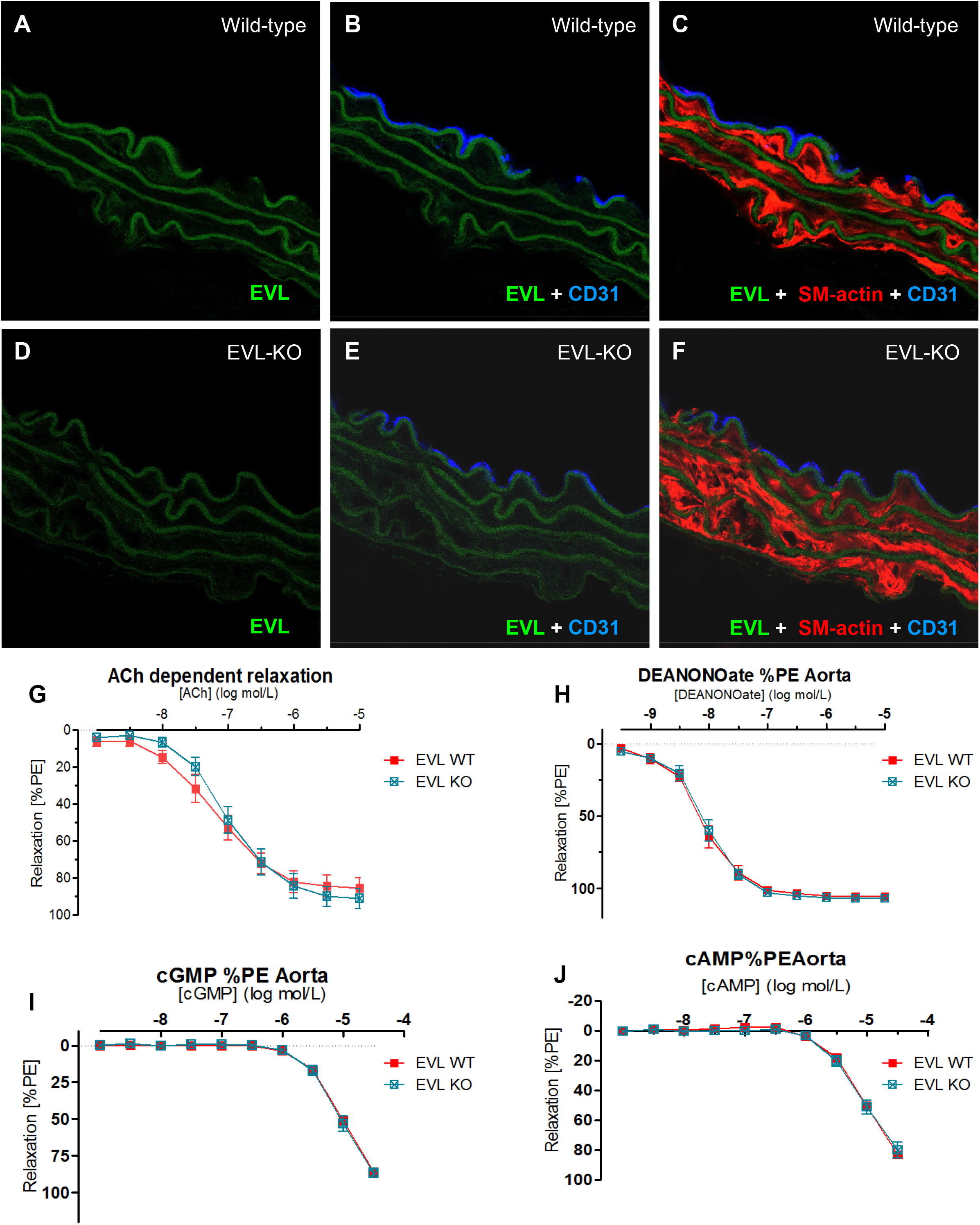
EVL-deficiency has no impact on the agonist-induced relaxation of aortic rings. (**A-F**) Confocal immunofluorescence microscopy of wild-type (**A-C**) and EVL-deficient (**D-F**) aortic rings stained with EVL-, CD31- and α-smooth muscle actin-specific antibodies. Representative images from 4 independent experiments are shown. (**G-J**) Myograph experiments with aortic rings from EVL^-/-^ mice or wild-type controls. Aortic rings were pre-contracted with 1 μM phenylephrin (PE) and then relaxed with increasing concentrations of acetylcholine (ACh), the NO-donor DeaNONOate, or the cAMP- or cGMP-analogs, Sp-5-6-DCI-BIMPS (cAMP) and 8-Br-pCPT-cGMP (cGMP), respectively. Data represent SEM ± SD, two-way ANOVA; N=6 animals, 4 rings per animal.

Little is known regarding the expression of Mena in vascular smooth muscle cells and there has in fact been some controversy about its presence in vascular smooth muscle cells. Indeed, while one group reported a robust expression of Mena expression in the vascular wall (Gambaryan et al., 2001), others were unable to reproduce this finding (Kim et al., 2010). To clarify these conflicting findings, we first analyzed the Mena promoter activity in smooth muscle tissue, making use of the Mena promoter-driven expression of the β-galactosidase reporter gene in Mena^GT/GT^ mice (Benz et al., 2013). X-gal staining of Mena^GT/GT^ tissue revealed a strong and specific Mena promoter activity in different types of smooth muscle, including the aorta and blood vessels of the heart, lung, kidney, liver and skeletal muscle, as well as in smooth muscle layers of stomach and intestine. Closer inspection of large vessels indicated that Mena promoter activity was predominant in the media. Mena promoter activity in the endothelium of vessels could not be confirmed by X-gal staining of cryosections (Figure 5).

**Figure 5.**
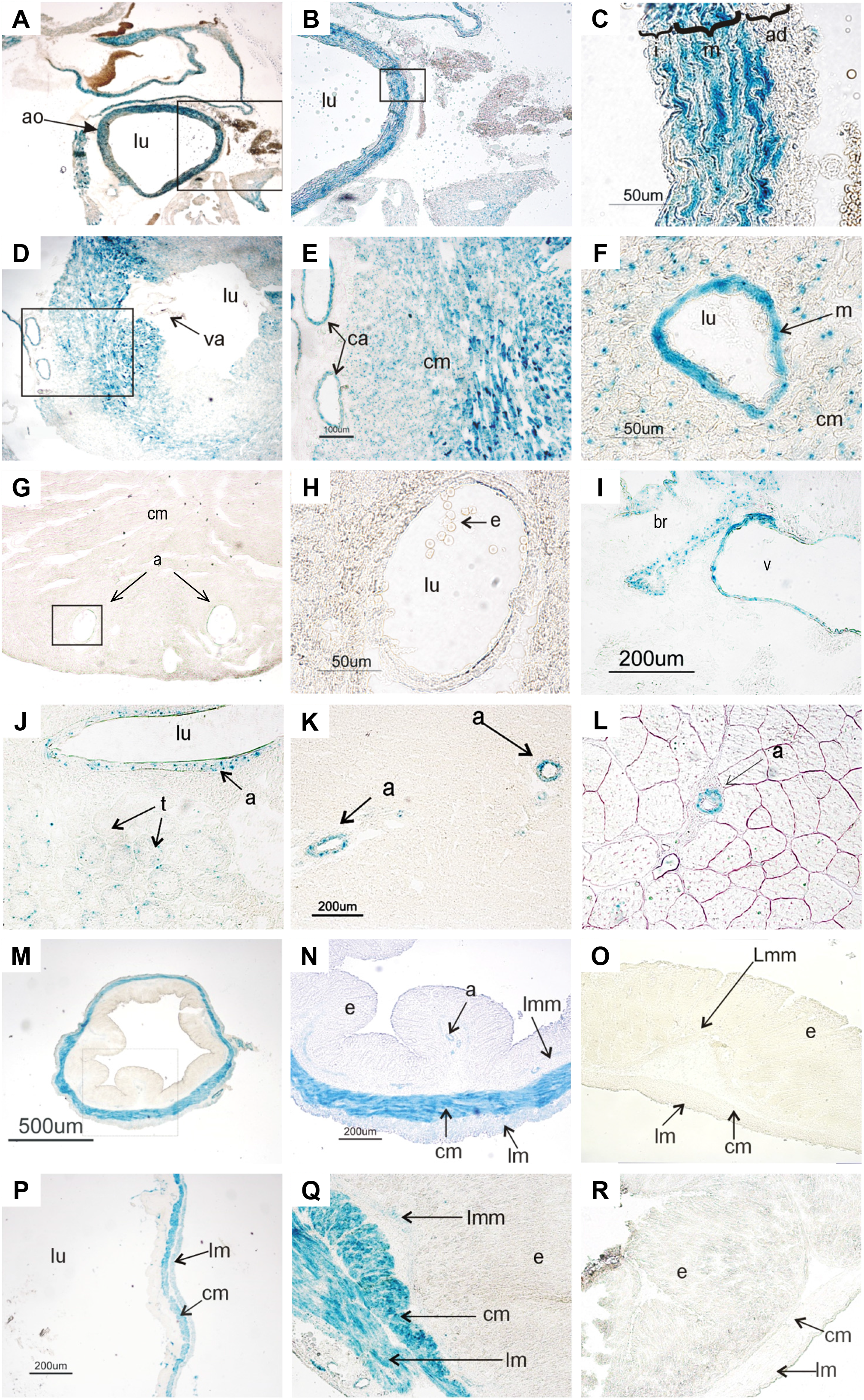
Mena promoter activity in vascular and gastrointestinal smooth muscle. X-gal staining of Mena^GT/GT^ cryosections and wild-type controls. Formation of blue indigo dye indicates β-galactosidase expression (Mena-promotor activity). (**A**) Mena^GT/GT^ mouse transversal section of the aorta, at the level of the heart base (ao, aorta; lu, lumen). (**B**) Enlargement of the indicated area in A; blue stained vessel wall (lu, lumen). (**C**) Enlargement of the indicated area in B; vessel wall layers intima (i), media (m), adventitia (ad) are indicated; blue staining predominantly observed in the media. (**D**) Mena^GT/GT^ mouse; transversal section of the left ventricle; vessels and heart muscle tissue are stained blue, valves (va) remain unstained (lu, lumen of ventricle). (**E**) Enlargement of the indicated area in D; walls of coronary arteries (ca) and cardiomyocytes (cm) are stained clearly. (**F**) Mena^GT/GT^ heart muscle tissue with coronary vessels (lu, lumen; m, media; cm, cardiomyocytes). (**G, H**) Wild-type mouse sections of the ventricular wall (negative control; cm, cardiomyocytes; a, arteries). (H) Enlargement of the indicated area in G showing branches of coronary arteries with erythrocytes (e) in the lumen (lu). (**I**) Mena^GT/GT^ mouse lung sections; walls of bronchioles (br) and vessels (v) show Mena promoter activity, alveolar tissue is not stained. (**J**) Mena^GT/GT^ sections of the kidney (renal cortex). Mena promotor activity is observed in renal artery (a) and in some tubules (t); (lu, lumen). (**K**) Mena^GT/GT^ liver-cryosection; Mena-promoter activity only detected in arteries (a). (**L**) Mena^GT/GT^ femoral skeletal muscle; β-galactosidase expression appears only in arteries (a) not in the muscle tissue. (**M**) Mena^GT/GT^ transversal section of the colon; intense blue colour indicates Mena promoter activity in the muscle layer. (**N**) Enlargement of the indicated area in M; showing blue stained arteries (a), lamina muscularis mucosae (lmm), circular muscle (cm) layer and longitudinal muscle (lm) layer; (e, epithelium). (**O**) Wild-type colon section; negative control (lmm, lamina muscularis mucosae; cm, circular muscle layer; lm, longitudinal muscle layer; e, epithelium). (**P**) Mena^GT/GT^ section of stomach wall (fundus/corpus area) showing Mena promoter activity in the muscle layer (cm, circular muscle layer; lm, longitudinal muscle layer; lu, lumen). (**Q**) Mena^GT/GT^ stomach wall (pylorus area; lmm, lamina muscularis mucosae; cm, circular muscle layer; lm, longitudinal muscle layer; e, epithelium). (**R**) Wild-type stomach section, negative control (cm, circular muscle layer; lm, longitudinal muscle layer; e, epithelium).

To target Mena gene expression *in vivo*, ES-cells were obtained from the European Conditional Mouse Mutagenesis Program (Enah^Gt(EUCE322f03)Hmgu^, parent cell line: E14TG2a, strain of origin: 129P2/OlaHsd) with the aim of generating global and tissue specific Mena knockout mice. Mena gene inactivation was mediated by the insertion of a conditional FlipRosaβgeo gene trap vector in intron 1 of the mouse Mena gene, upstream of exons coding for the N-terminal EVH1 domain. Due to a splice acceptor and a polyA signal in the vector, Mena transcripts were prematurely terminated resulting in a Mena protein null mutant. The mutation could be reversed and re-introduced through two site-specific recombination systems, FLPe_frt/F3 and Cre_loxP/lox511, respectively, that enable gene trap cassette inversions from the sense, coding strand of a trapped gene to the antisense, noncoding strand and back (Schnutgen et al., 2005) (Figure 6A). ES cells were used to generate chimeras by injections of C57BL/6 host blastocysts and germline transmission of the disrupted allele was verified by PCR and splinkerette-PCR. In contrast to a previous publication (Lanier et al., 1999), bi-allelic deletion of Mena in homozygous mutant animals (Mena^F03/F03^) was lethal and they died in utero. On embryonic day (E) 11 Mena^F03/F03^ embryos were small and runted and displayed craniofacial defects reminiscent of Mena/VASP-double deficient mice (Menzies et al., 2004) (Figure 6B-D). Currently, it remains unclear why Mena deletion induced such different phenotypes. Potential explanations include incomplete Mena gene inactivation, e.g., due to splicing events, or the varying genetic background of the mutant mice. To rescue the lethal phenotype and generate smooth-muscle-specific Mena^-/-^ mice, we first inverted the βgeo-cassette by mating with a FLPe-deleter strain (Zink et al., 2021), generating conditional Mena alleles (Mena^INV/INV^); and subsequently with mice expressing the Cre recombinase under the control of the smooth muscle myosin heavy chain promoter (Groneberg et al., 2010). After Tamoxifen treatment, a time-dependent reduction of Mena protein expression in the aorta and the mesenteric artery was observed. While Mena protein was still readily detectable one week after tamoxifen treatment, Mena protein levels were markedly reduced after six weeks, as detected by Western blotting (Figure 7A). However, it seems that the turnover of the Mena protein is slow and that, similar to the case of the soluble guanylyl cyclase (sGC) (Groneberg et al., 2010), an even longer time after tamoxifen treatment may be required to fully deplete the Mena protein from smooth muscle cells. Consistent with our promoter studies, confocal microscopy of wild-type aortas revealed high Mena levels in smooth muscle cells, but Mena protein expression in endothelial cells was hardly detectable (Figure 7B-G).

**Fig. 6.**
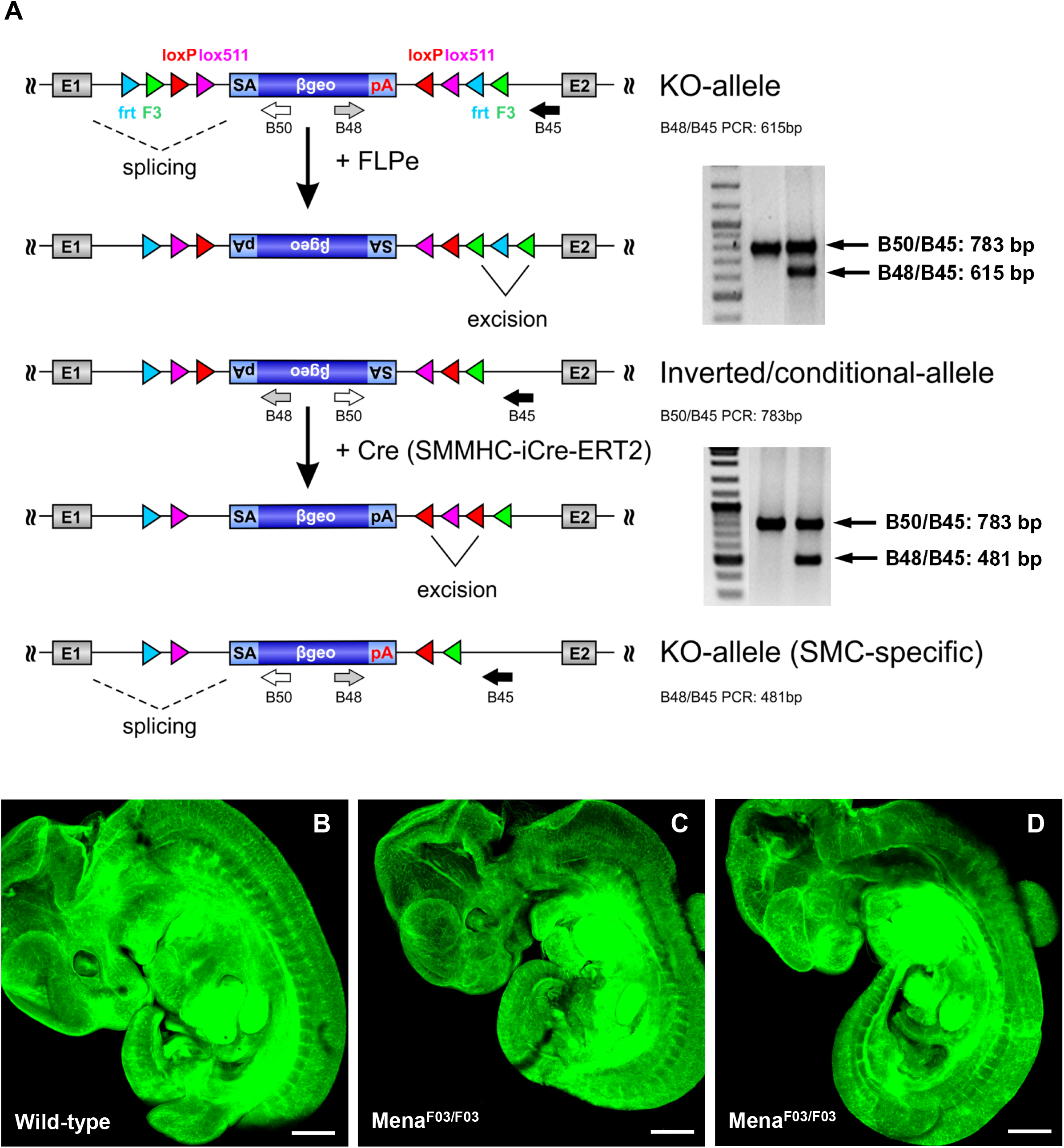
Generation and lethal phenotype of MenaF^03/F03^ knockout mice. (**A**) Conditional gene trap vector and Mena gene inactivation in Enah^Gt(EUCE322f03)Hmgu^ mice. Mena gene inactivation is mediated by the insertion of a conditional FlipRosaβgeo gene trap vector in intron 1 of the mouse Mena gene, upstream of exons coding for the N-terminal EVH1 domain. Due to a splice acceptor (SA) and a polyA (pA) signal in the vector, Mena transcripts are prematurely terminated resulting in a Mena protein null mutant and instead, expression of the β-geo gene, a fusion of β-galactosidase and neomycin resistance. The mutation can be reversed and re-introduced through two site-specific recombination systems, FLPe_frt/F3 and Cre_loxP/lox511, respectively, that enable gene trap cassette inversions from the sense coding strand of a trapped gene to the antisense noncoding strand and back. The status and orientation of the gene trap cassette can be monitored by PCR, yielding a 615bp B48/B45 PCR product for the original KO-allele, a 783bp B50/B45 PCR product for the inverted/conditional allele, or a 481bp B48/B45 PCR product for the re-inverted KO allele. (**B-D**) Homozygous Mena^F03/F03^ mice are lethal and die in utero. E11 Mena^F03/F03^ embryos or wild-type littermates were fixed, cleared and stained with CD31-specific antibodies to visualize blood vessels. Maximum projections of confocal Z-stack images are shown for one wild-type and two Mena^F03/F03^ embryos. E11 Mena^F03/F03^ embryos were small and runted and displayed craniofacial defects.

**Figure 7.**
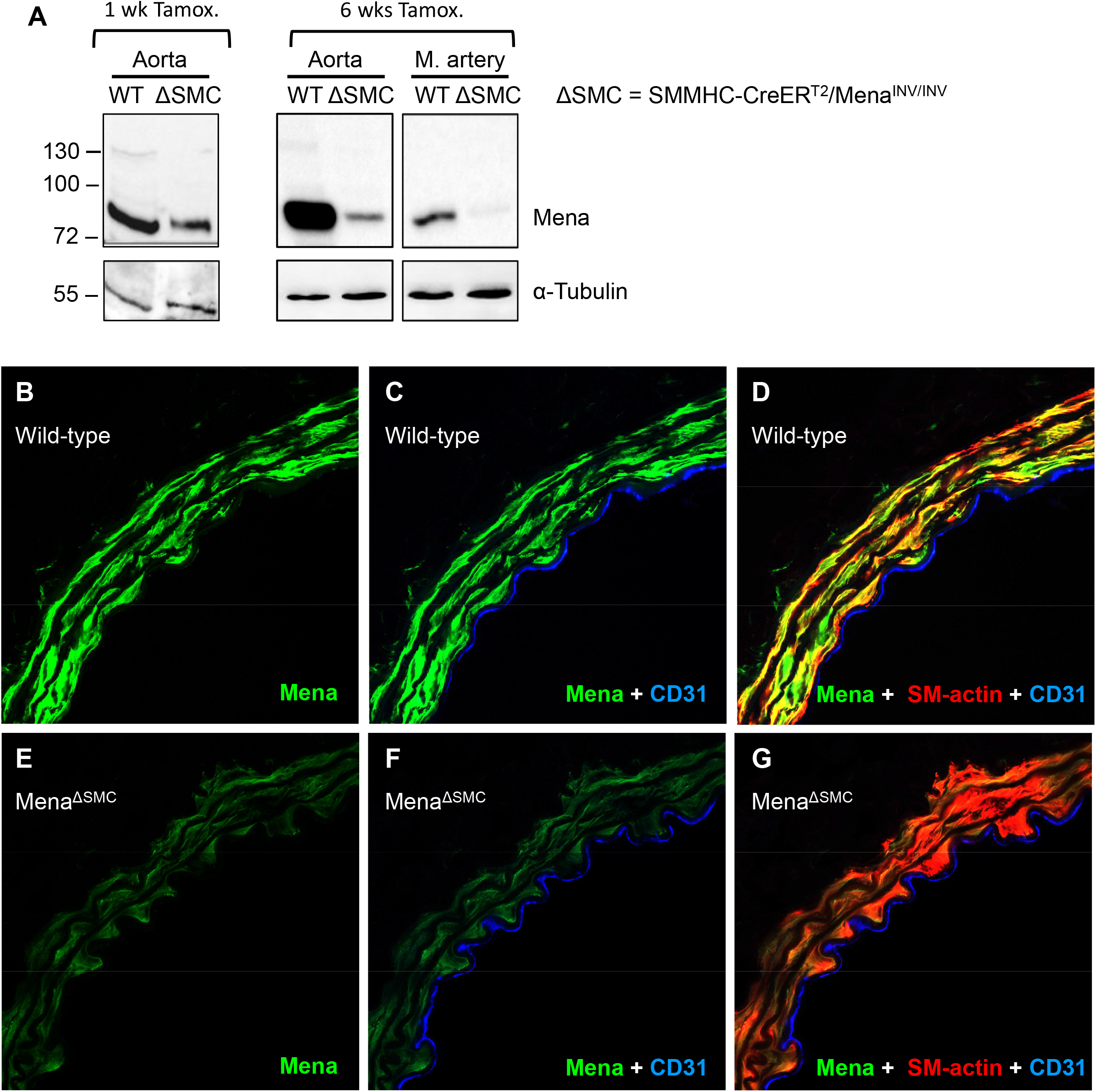
Smooth muscle specific Mena-deletion (Mena^ΔSMC^). (**A**) Western blots showing the time-dependent reduction of Mena protein expression in the aorta and the mesenteric artery of Mena^ΔSMC^ mice, 1 and 6 weeks after Tamoxifen treatment. (**B-G**) Confocal microscopy demonstrating reduced Mena expression in aortic smooth muscle of Mena^ΔSMC^ mice, 6 weeks after Tamoxifen treatment. Staining with CD31- and α-smooth muscle actin-specific antibodies was used to identify endothelial cells and smooth muscle cells, respectively. Representative images from 4 independent experiments are shown.

Given the strong expression of Mena and VASP in vascular smooth muscle and the fact, that the preferred PKA and PKG phosphorylation sites are conserved in the two proteins (Figure 1), we investigated how combined Mena/VASP deletion would impact on the agonist-induced smooth muscle relaxation. We performed organ bath experiments with aortic rings of VASP^-/-^Mena^GT/GT^ mice that completely lack VASP expression and display only minimal Mena protein levels in cardiovascular cells, including blood vessels (Benz et al., 2013). In comparison to their wild-type controls, aortic rings from VASP^-/-^Mena^GT/GT^ mice displayed a reduced responsiveness to NO and smooth muscle relaxation to low concentrations (up to 0.1 μM) of DEA-NO was significantly impaired (Figure 8A). Further support that the NO/cGMP/PKG and the cAMP/PKA cascade induces lower relaxation in VASP^-/-^Mena^GT/GT^ mice was obtained by experiments targeting the PKG and PKA directly. Therefore, after pre-contraction with phenylephrine, the cAMP-analog (Sp-5-6-DCI-BIMPS; 10 μM) and cGMP-analog (8-Br-pCPT-cGMP; 10 μM) were applied to relax the aortic rings. In rings from VASP^-/-^Mena^GT/GT^ mice, cGMP- and cAMP-induced relaxation was more than 50% reduced as compared to wild-type controls, indicating that PKG- and PKA-dependent smooth muscle relaxation is significantly disturbed in the absence of Mena and VASP (Figure 8B-C).

**Figure 8.**
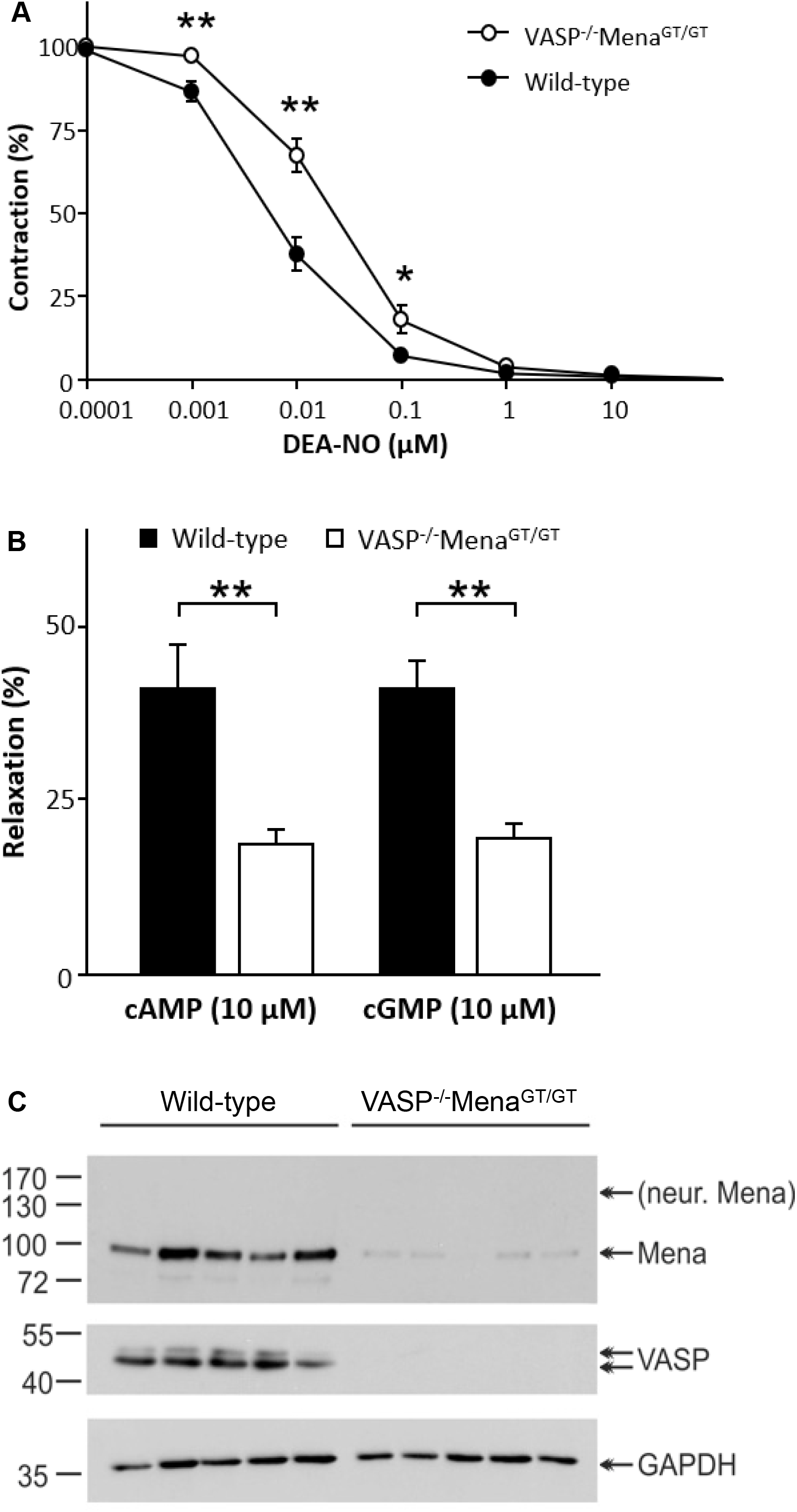
Impact of combined Mena/VASP deletion on aortic smooth muscle relaxation. (**A, B**) Myograph experiments with aortic rings from VASP^-/-^Mena^GT/GT^ or wild-type controls. Aortic rings were pre-contracted with 1 μM phenylephrin (PE) and then relaxed with increasing concentrations of the NO-donor DeaNONOate (DEA-NO), or the cAMP- or cGMP-analogs Sp-5-6-DCI-BIMPS (cAMP) and 8-Br-pCPT-cGMP (cGMP), respectively. (N=5 animals per group, two rings per animal; \**P*<0.05, \*\**P*<0.01). After the experiments, Mena and VASP expression in the rings was analyzed by Western blotting (**C**).

## Discussion

In the present study, we investigated the impact of individual and combined deletion of Ena/VASP proteins on vascular smooth muscle relaxation. In contrast to previous studies (Aszodi et al., 1999), we found that VASP deficiency significantly impaired the ACh-induced and NO-dependent relaxation of aortic rings *ex vivo*. However, this effect was age-dependent and only observed in aortic rings from 7-month-old animals, and not in tissues from 3-month-old animals. Interestingly, ACh-induced and NO-dependent relaxation was already impaired in mesenteric artery rings from 4-month-old VASP^-/-^ mice, indicating that resistance arteries/arterioles may be more susceptible to VASP-deficiency. The latter fits well with the recent report of impaired NO-mediated dilation in the microcirculation of VASP-deficient mice (Poley et al., 2023), indicating that VASP is also important for dilation of vascular smooth muscle *in vivo*.

In contrast to VASP, EVL was not detectable in the wall of the aorta and EVL-deficiency had no impact on smooth muscle relaxation. This indicates a more specialized function of EVL in different vascular beds. Indeed, EVL was previously implicated in endothelial barrier function and sprouting angiogenesis (Mascarenhas et al., 2022; Zink et al., 2021), indicating a potential role in non-quiescent endothelial cells in capillaries. This contrasting function is reflected by the fact that EVL lacks the preferred PKG-dependent phosphorylation site, which is conserved in VASP and Mena, perhaps indicating a critical role of this serine in regulating vascular tone. Mena on the other hand, was strongly expressed in various smooth muscle tissues, including blood vessels. We therefore hypothesized that Mena could at least partially compensate the loss of VASP in smooth muscle relaxation and analyzed aortic rings of VASP^-/-^Mena^GT/GT^ mice, which lack VASP completely and express only minimal levels of Mena in the aorta. Consistent with a redundant role of Mena and VASP, NO-mediated relaxation of aortic rings from VASP^-/-^Mena^GT/GT^ mice were more severely impaired than those from VASP^-/-^ mice. Moreover, cAMP- and cGMP-induced relaxations were also significantly impaired in aortic rings from VASP^-/-^ Mena^GT/GT^ mice, which was not the case in mice that lacked only VASP. However, additional head-to-head experiments with complete Mena knockout tissue are required to more clearly decipher the individual contribution and functional redundancy of the two proteins.

The molecular mechanisms underlying Mena/VASP-dependent smooth muscle cell relaxation is currently elusive. Given that ACh-induced, NO-dependent but not cGMP-dependent smooth muscle relaxations were impaired in VASP^-/-^ vessels, this suggests that regulation of cGMP synthesis, turnover or compartmentalization may be impaired in VASP-deficient animals. Indeed, a VASP-sGC feedback loop was previously proposed to exist in adipocytes (Jennissen et al., 2012). However, sGC and cGMP levels were actually *increased* in VASP-deficient adipose tissue and further studies are required to elucidate a potential feedback loop in vascular smooth muscle tissue.

Apart from the regulation of sGC/cGMP signaling, the molecular mechanism(s) underlying impaired smooth muscle relaxation in VASP and/or Mena-deficient vessels could also be related to actin dynamics. In fact, ample evidence supports an important contribution of actin and actin binding proteins for regulating the development of mechanical tension in smooth muscle cells. The functional role of actin polymerization during contraction is likely independent from the Ca^2+^ triggered actomyosin crossbridge cycle (Gunst and Zhang, 2008). Contractile stimulation is thought to initiate the assembly of cytoskeletal/extracellular matrix adhesions and actin polymerization at the cortex, which strengthen the membrane for the transmission of force generated by the contractile machinery (Gunst and Zhang, 2008; Tang, 2015; Yamin and Morgan, 2012). Conversely, inhibition of actin polymerization may serve to relax smooth muscle tissue. In vascular smooth muscle cells, VASP is located in dense plaques and dense bodies (Markert et al., 1996), in which actin filaments are anchored to the extracellular matrix and within the cytosol, respectively (Gunst and Zhang, 2008). In large arteries, α-smooth muscle actin is the major constituent of thin filaments, but β- and γ-cytoplasmic actin and γ-smooth muscle actin isoforms are also present. Interestingly, β-cytoplasmic actin is associated with dense plaques and dense bodies (Yamin and Morgan, 2012), suggesting that VASP is a selective regulator of β-cytoplasmic actin dynamics in vascular smooth muscle cells. This is strongly reminiscent of a previous study in heart muscle tissue, where Mena specifically colocalized with the β-cytoplasmic actin isoform, but not with α-cardiac or γ-cytoplasmic actin fibers (Benz et al., 2013). VASP was shown to co-localize with hot spots of actin polymerization at the cell cortex of vascular smooth muscle cells and VASP phosphorylation, known to inhibit actin polymerization, was decreased after stimulation with the contractile agent phenylephrine (Kim et al., 2010). In airway smooth muscle, contractile stimulation with acetylcholine triggered the PKC-mediated phosphorylation of VASP at S157 and the formation of VASP-vinculin complexes at membrane adhesion sites, which is a necessary for VASP-mediated actin polymerization and tension generation. Although forskolin, which induces cAMP/PKA signaling, also induced VASP S157 phosphorylation and membrane localization, it did not stimulate actin polymerization, potentially because forskolin also strongly induced the S239 phosphorylation of the protein (Wu and Gunst, 2015), which is known to inhibit actin polymerization (Benz et al., 2009). This observation may therefore constitute a potential mechanism for cAMP/PKA- and cGMP/PKG-induced smooth muscle relaxation mediated by VASP (and Mena).

In summary, our study has revealed an important function of Mena and VASP in the agonist-induced vascular smooth muscle relaxation, but further studies are required to clarify the underlying molecular mechanisms and the relevance of Ena/VASP proteins for blood pressure control *in vivo*.

## Materials and Methods

### Mouse strains

VASP^-/-^, EVL^-/-^, Mena^GT/GT^, VASP^-/-^Mena^GT/GT^, FLPe-deleter, and mice expressing Cre recombinase under the control of the smooth muscle myosin heavy chain promoter were reported previously (Benz et al., 2013; Groneberg et al., 2010; Hauser et al., 1999; Zink et al., 2021). To generate Mena(F03) mice, ES-cells from the European Conditional Mouse Mutagenesis Program (Enah^Gt(EUCE322f03)Hmgu^, parent cell line: E14TG2a, strain of origin: 129P2/OlaHsd) used to generate chimeras by injections of C57BL/6 host blastocysts and germline transmission of the disrupted allele was verified by PCR and splinkerette-PCR. Status/orientation of the gene trap cassette was monitored by PCR, yielding a 615bp B48/B45 PCR product for the original KO-allele, a 783bp B50/B45 PCR product for the inverted/conditional allele, or a 481bp B48/B45 PCR product for the re-inverted KO allele. B045: 5′-CTCCGCCTCCTCTTCCTCCATC-3′; B048: 5′-TCCCACTGTCCTTTCCTAATAA-3′, B050: 5′-TTTGAGGGGACGACGACAGTAT-3′.

### Immunohistochemistry

Ten micrometer cryosections of aorta and mesenteric artery were stained with standard methods. Briefly, sections were fixed in 4% formalin on ice, permeabilized with 0.2% TX-100 in PBS and blocked with 10% normal donkey serum in PBS. Primary antibodies: anti-Mena (rabbit, 1:600; (Benz et al., 2013)), anti-VASP (rabbit, 1:150, (Benz et al., 2008)), anti-EVL (rabbit, 1:80, (Zink et al., 2021)), anti-CD31 (rat, BD Pharmingen 553370, 1:200), anti-smooth muscle actin (mouse, Sigma-Aldrich Sigma Aldrich C6198, 1:500); secondary antibodies: Alexa Fluor conjugated donkey anti-mouse/rabbit/rat antibodies, 1:300. Sections were imaged on a Leica SP8 microscope equipped with a 60x immersion objective.

### Tissue clearing and 3D imaging of embryonic day 11 wild-type and Mena^F03/F03^ embryos

Dissected embryos were fixed overnight in Roti™Histofix (Carl Roth, Carlsruhe, Germany) at 4°C and then permeabilized and blocked overnight (37°C) with 0.5% Triton X-100, 5% donkey serum (Merck, Darmstadt, Germany) and 1% DMSO in PBS. Embryos were then stained with primary CD31 (clone MEC13.3, BD Bioscience, Heidelberg, Germany) and secondary antibodies overnight in the same buffer. Tissue clearing was performed as previously described (Erturk et al., 2012). Embryos were dehydrated in one hour intervals in an ascending tetrahydrofuran (THF, Merck, Darmstadt, Germany) series (50% THF, 70% THF, 80% THF, and 100% THF) at room temperature. After a final de-lipidation with Dichloromethane (Merck, Darmstadt, Germany) for 15 minutes, embryos were transferred to dibenzylether (Merck, Darmstadt, Germany) and imaged after a final incubation step of at least 3 hours. Z-stacks were acquired with a Leica SP8 microscope equipped with a 5x objective and the Leica application suite X was used to generate maximum projection images of the embryos.

### Organ Bath Experiments

Thoracic aortas and mesenteric arteries were cut into rings and mounted in a myograph (DMT model 610M or 620M) as previously described (Groneberg et al., 2010) with a resting tension of 5 or 9.81 mN in the present of diclofenac (10 μM). After equilibration (at least 60 minutes at 37°C), rings were precontracted with phenylephrine (1 μM; Sigma-Aldrich P6126). To investigate endothelial nitric oxide synthase (eNOS)-dependent smooth muscle relaxation, rings were relaxed with increasing concentrations of acetylcholine (ACh; Sigma-Aldrich A6625). eNOS-independent relaxation was tested in presence of the pharmacologic eNOS inhibitor N-nitro-L-arginine methyl ester (L-NAME, 300 μM; Sigma-Aldrich N5751) either using increasing concentrations of the NO-donor dimethylamine nonoate (DEA-NONOate; Sigma Aldrich D184) or the stable cGMP analog 8-Br-pCPT-cGMP (cGMP; BioLog C009 E) to directly activate PKG. Similarly, direct PKA activation was achieved using increasing concentrations of the stable cAMP analog Sp-5,6-DCl-cBIMPS (cAMP; BioLog D014). Isobutyl methylxanthine (IBMX; Sigma-Aldrich 15879) was used to induce maximal relaxation. Each experiment was performed in parallel with 2 to 4 rings.

### X-Gal staining of Mena^GT/GT^ cryosections

Organ cryosections of Mena^GT/GT^ mice with wild-type sections as negative controls were used. The following reagents were prepared and used as stock solutions: 0.1M phosphatase buffer (3.75 g NaH_2_ PO_4_-H_2_O, 10.35 g Na_2_HPO_4_ dissolved in 1 liter ddH_2_O pH 7.3); fix buffer for X-gal staining (0.1 M phosphatase buffer (pH 7.3), 5 mM EGTA, 2 mM MgCl_2_, 0.2% glutaraldehyde); wash buffer for X-gal staining (0.1 M phosphate buffer (pH 7.3), 2 mM MgCl_2_); X-gal staining buffer (0.1 M phosphate buffer (pH 7.3), 2 mM MgCl_2_, 5 mM K_4_Fe(CN)_6_ -3H_2_0, 5 mM K_3_Fe(CN)_6_; directly before use: X-gal (5-bromo-4-chloro-3-indolyl-β-D-galactoside) was added to a final concentration of 1 mg/ml); X-gal stock solution (50 mg/ml) (10 ml dimethylformamide, 500 mg X-gal), Mowiol + Dabco (2.4 g Mowiol 4-88 (Hoechst) 6 g Glycerol, 6 ml H_2_O, 12 ml 0.2M Tris (pH 8.5), 2.5% 1.4-diazobicyclo-octane).

Approximately 2 - 4 hours before staining, X-gal solution was added to X-gal staining buffer to a final concentration of 1 mg/ml and mixed thoroughly. Prior to use, this solution was filtered with a folding filter in order to remove any precipitate. Straight after the collection of the cryosections, the object slides were transferred to a glass chamber with fix buffer. After 15 min of fixation, the object slides were washed twice with wash buffer, each time for 5 min. Finally, the cryosections were covered with the filtered X-gal staining buffer, and incubated for 13 to 15 hours at 37°C. The next day, object slides were washed twice, and then mounted with 100 μl of Mowiol and a cover slip. After air-drying at room temperature overnight, stained sections were stored at 4°C until inspection using a Nikon microscope. Images were acquired using the ACT-1 software (Nikon).

## Acknowledgements

This work was supported by the Deutsche Forschungsgemeinschaft (SFB 834/A8 to PMB, SFB 834/A5 to IF). PMB was also supported by the German Center for Cardiovascular Research (DZHK B14-028 SE). The authors acknowledge Stefan Offermanns and Nina Wettschureck (Planck Institute for Heart and Lung Research, Bad Nauheim, Germany) for help with the generation of Mena knockout mice, Bernhard Nieswandt (University of Würzburg, Germany) for providing transgenic FLP-deleter mice, and Andreas Friebe (University of Würzburg, Germany) for scientific input.

